# TAS1R3 regulates GTPase signaling in human skeletal muscle cells for glucose uptake

**DOI:** 10.1101/2025.09.10.674099

**Authors:** Joseph M. Hoolachan, Rekha Balakrishnan, Karla E. Merz, Debbie C. Thurmond, Rajakrishnan Veluthakal

**Author notes:** <u>Corresponding authors</u>: Debbie C. Thurmond, Ph.D, Rajakrishnan Veluthakal, Ph.D. Contributed equally (Joseph M. Hoolachan, Rekha Balakrishnan).

## Abstract

**Background:** Taste receptor type 1 member 3 (TAS1R3) is a class C G protein-coupled receptor (GPCR) traditionally associated with taste perception. While its role in insulin secretion is established, its contribution to skeletal muscle glucose uptake a process responsible for 70–80% of postprandial glucose disposal remains unclear.

**Methods:** TAS1R3 expression was assessed in skeletal muscle biopsies from non-diabetic and type 2 diabetes (T2D) donors using qPCR and immunoblotting. Functional studies in human LHCN-M2 myotubes involved TAS1R3 inhibition with lactisole or siRNA-mediated knockdown, followed by measurement of insulin-stimulated glucose uptake using radiolabeled glucose assays. Rac1 activation and phospho-cofilin were analyzed by G-LISA and Western blotting, and Galphaq/11 involvement was tested using YM-254890.

**Results:** TAS1R3 mRNA and protein levels were significantly reduced in T2D skeletal muscle. Pharmacological inhibition or knockdown of TAS1R3 impaired insulin-stimulated glucose uptake in myotubes.

**Conclusion:** TAS1R3 regulates skeletal muscle glucose uptake through a non-canonical insulin signaling pathway involving Rac1 and phospho-cofilin, independent of IRS1-AKT and Galphaq/11 signaling. These findings identify TAS1R3 as a key determinant of Rac1-mediated glucose uptake and a potential therapeutic target for improving insulin sensitivity in T2D.

## Background

Over 750 million people globally have prediabetes. Without intervention, 50% of individuals with prediabetes will progress to type 2 diabetes (T2D) annually (1). In 2021, the global age-adjusted prevalence of impaired glucose tolerance (IGT) among adults aged 20–79 years was estimated at 9.1%, corresponding to approximately 464 million individuals. This prevalence is projected to rise to 10.0% (∼ 638 million individuals) by 2045. Similarly, in 2021, an estimated 5.8% of adults (20-79 years) (∼ 298 million people) had impaired fasting glucose (IFG). By 2045, this figure is expected to increase to 6.5%, affecting roughly 414 million individuals (2). Skeletal muscle insulin resistance is a major pathological feature in T2D (3). Peripheral insulin resistance is a key driver of prediabetes and its progression into T2D, with skeletal muscle being a major target (3–5). In healthy individuals, skeletal muscle plays a central role in maintaining whole-body glucose homeostasis, accounting for ∼70-90% of insulin-stimulated peripheral blood glucose uptake (6–8). However, prolonged skeletal muscle insulin resistance can drive myopathic comorbidities in T2D patients including progressive muscle loss (sarcopenia) (9), exercise intolerance (10, 11), skeletal muscle inflammation (12) and impaired muscle repair (13). Weight management is a key intervention for mitigating insulin resistance and T2D, and the widely used glucagon-like peptide-1 receptor agonists (GLP-1 RAs) are associated with an elevated risk of sarcopenia (14–17). Therefore, novel therapeutic strategies that can effectively prevent or reverse skeletal muscle insulin resistance, albeit without any adverse myopathic risks are needed.

Although multiple factors are attributed to the development of skeletal muscle insulin resistance including defective glucose and lipid utilization (18), progressive mitochondrial decline (19), aging (20) and oxidative stress (21), a key feature is the impairment of the canonical (PI3K/AKT) (22, 23) and non-canonical (GTPase/Actin remodeling) insulin signaling cascades (24). Normally, both cascades require stimulation of the insulin receptors; however, the non-canonical insulin signaling pathway is able to stimulate GLUT4-mediated vesicle translocation to the sarcolemma independent of Insulin receptor substrate 1 (IRS1) activation (25). Indeed, several G-protein coupled receptor pathways are able to activate the small Rho-family GTPase Rac1 to promote actin filament remodeling to promote GLUT4 translocation (26). Although this provides an insulin-independent source for GLUT4 translocation, the upstream regulators that coordinate small GTPase activity in skeletal muscle remain incompletely defined.

One potential candidate to coordinate small GTPase activity in skeletal muscle is the sweet taste receptor TAS1R3, a G protein-coupled receptor (GPCR); TAS1R3 knockout mice show impaired glucose tolerance (27). Recent studies have demonstrated that TAS1R3/TAS1R2 heterodimer is actively expressed in skeletal muscle tissues (28). This suggests a potential non-gustatory role for TAS1R in skeletal muscle physiology, possibly linked to nutrient sensing, energy metabolism, or insulin signaling. Given that a number of conserved molecular pathways play a role in both β-cell GSIS and skeletal muscle insulin-stimulated glucose uptake (29–34), TAS1R3 could also function in skeletal muscle glucose uptake.

In this study, we hypothesize that TAS1R3 is required for insulin-stimulated glucose uptake in skeletal muscle *via* Rac1-mediated non-canonical signaling. Using T2D and non-diabetic human skeletal muscle biopsies, as well as clonal human LHCN-M2 myotube cell lines, we aim to understand the significance of TAS1R3 signaling in skeletal muscle insulin resistance.

## RESULTS

### TAS1R3 transcript and protein levels are decreased in T2D human skeletal muscle

To investigate the impact of diabetogenic stress on TAS1R3 expression, we assessed TAS1R3 mRNA levels in human LHCN-M2 myotubes exposed for 16 h to glucolipotoxic (GLT) conditions (25 mM glucose and 200 μM palmitic acid). TAS1R3 transcript levels were significantly reduced in human LHCN-M2 myotubes post-GLT exposure (**Fig. 1A**). In parallel, insulin-stimulated glucose uptake was markedly diminished in these cells (**Fig. 1B**), further indicating the onset of insulin resistance. These findings are consistent with previous reports demonstrating that palmitate exposure induces insulin resistance, thereby reducing glucose uptake in L6 skeletal muscle myotubes and in human primary myotubes (35, 36). Next, we assessed TAS1R3 levels in T2D and non-diabetic (ND) human skeletal muscle biopsies (**Table 1**). Both TAS1R3 mRNA levels and protein abundances were significantly reduced (> 50%) in T2D skeletal muscle versus non-diabetic biopsies (**Fig. 1C-D**). Thus, TAS1R3 deficiency may contribute to peripheral glucose uptake dysfunction in T2D skeletal muscle.

**Figure 1.**
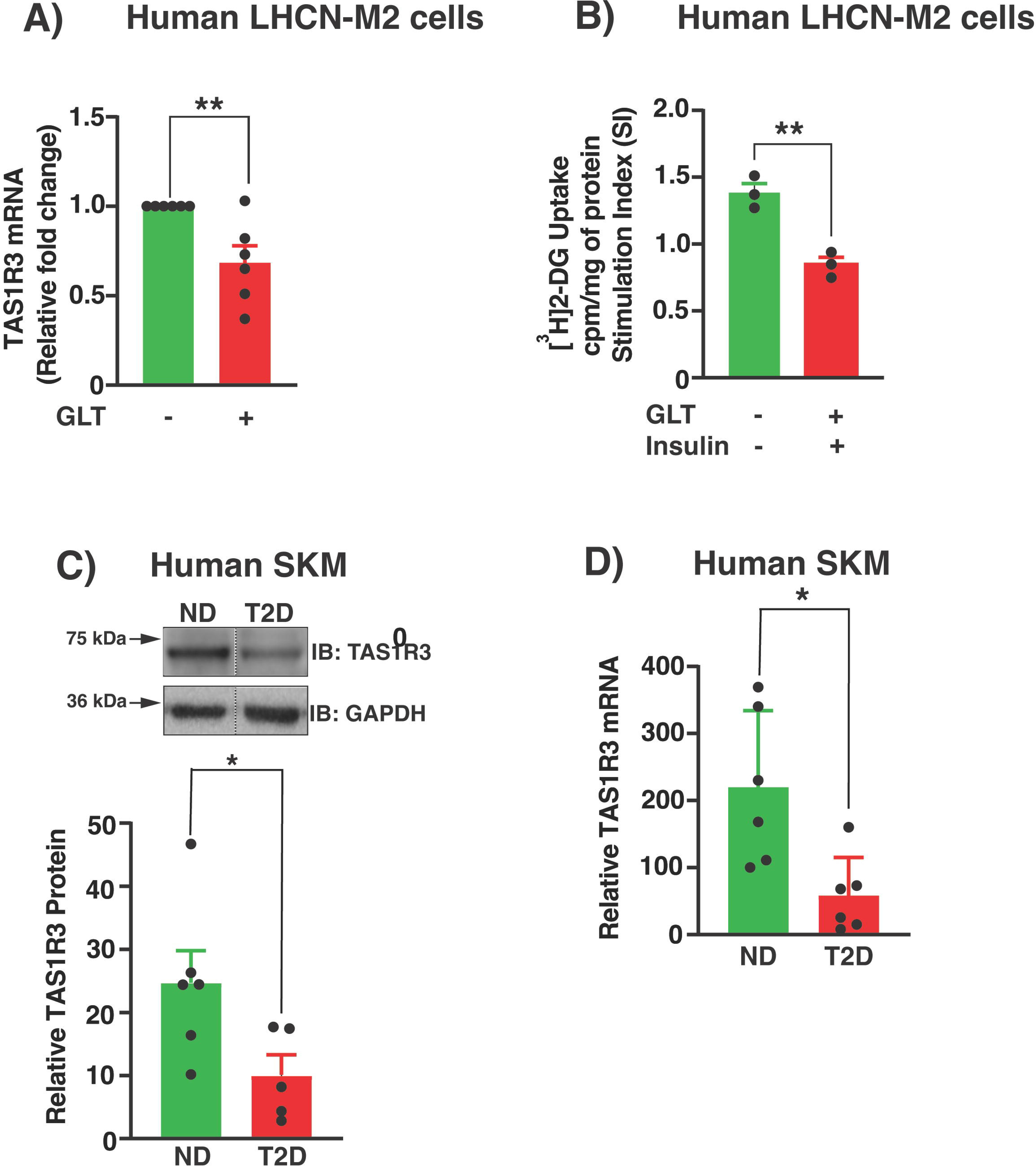
Human T2D skeletal muscle has lower TAS1R3 abundance. (**A-B**) LHCN-M2 myotube cells were untreated or exposed to GLT stress for up to 16 h. (A) TAS1R3 mRNA was expressed relative to tubulin mRNA. (n=6 independent experiments). (**B**) Glucose ([^3^H]2-DG) uptake ± insulin stimulation was normalized to protein content. Cpm: Counts per minute (n=3 independent experiments). Stimulation index (SI) was calculated as glucose uptake with insulin divided by the glucose uptake at basal. (**C**) Densitometry analysis of TAS1R3 protein in T2D (n=5 donors) versus ND (n=6 donors) skeletal muscle. Top: representative immunoblot (IB). TAS1R3 protein was expressed as relative to GAPDH. Vertical dashed lines indicate splicing of lanes from within the same gel exposure. (**D**) Quantification of TAS1R3 mRNA expression in human T2D (n=6 donors) versus ND (n=6 donors) skeletal muscle using qPCR. TAS1R3 mRNA was expressed as relative to tubulin mRNA. **(A-D**). Data expressed as mean ± SEM. *p < 0.05, **p < 0.01.

**Table 1:** Non-diabetic and T2D human skeletal muscle donor profiles.

| **NDRI #** | **Age** | **Race** | **Gender** | **BMI** | **Condition** |
| --- | --- | --- | --- | --- | --- |
| ND09743 | 46 | Caucasian | M | 34.9 | ND |
| ND09744 | 62 | Caucasian | M | 28.2 | ND |
| ND09749 | 49 | Caucasian | F | 21.6 | ND |
| ND09706 | 45 | Caucasian | M | 44.5 | T2D |
| ND09754 | 67 | Caucasian | M | 32.3 | T2D |
| ND10105 | 53 | Caucasian | F | 28 | T2D |
| ND13347 | 78 | Caucasian | M | 26.4 | ND |
| ND13230 | 68 | Caucasian | F | 40.2 | ND |
| ND13216 | 54 | Caucasian | M | 31.3 | ND |
| ND13214 | 74 | Caucasian | M | 32.1 | T2D |
| ND13363 | 59 | Caucasian | M | 41.5 | T2D |
| ND13239 | 58 | Caucasian | M | 50.3 | T2D |

ND= Non-diabetic individuals; T2D= Type 2 diabetic individuals; BMI= Body mass index;
M= Male, F= Female; NDRI# = National disease research interchange (NDRI) sample number, provided muscle source as quadriceps or 'leg muscle'.

### TAS1R3 inhibition impedes insulin-stimulated glucose uptake via a non-canonical insulin signaling pathway

To determine the requirement for TAS1R3 in skeletal muscle peripheral glucose uptake, we evaluated the impact of TAS1R3 (Lactisole; TAS1R3*(i)*) or Gαq/11 (YM-25490; Gαq/11(i)) inhibition against a vehicle control in insulin-stimulated human LHCN-M2 myotubes by measuring 2-deoxyglucose ([^3^H]2-DG) uptake. Skeletal muscle is a major site for glucose disposal and plays a critical role in systemic glucose homeostasis. Previous studies have shown that skeletal muscle–specific activation of Gq signaling improves glucose uptake and insulin sensitivity, even under metabolic stress (37). Additionally, CaSR, a Gαq/11-coupled receptor activated by kokumi substances, enhances sweet, salty, and umami perception *via* TAS1R3 components (38), highlighting Gαq/11 signaling as a key integrator of nutrient sensing and metabolic regulation across tissues. This overlap suggests that Gαq/11 signaling integrates nutrient sensing and metabolic regulation across multiple tissues. Therefore, investigating Gαq/11 in skeletal muscle provides insight into whether TAS1R3 couples Gq mechanisms to downstream signaling components to glucose uptake. TAS1R3 inhibition significantly impeded glucose uptake by ∼50% versus the vehicle in insulin-stimulated human LHCN-M2 myotubes (**Fig. 2A**). We also observed that TAS1R3 inhibition reduced glucose uptake under basal conditions (**Fig. 2A**).

**Figure 2.**
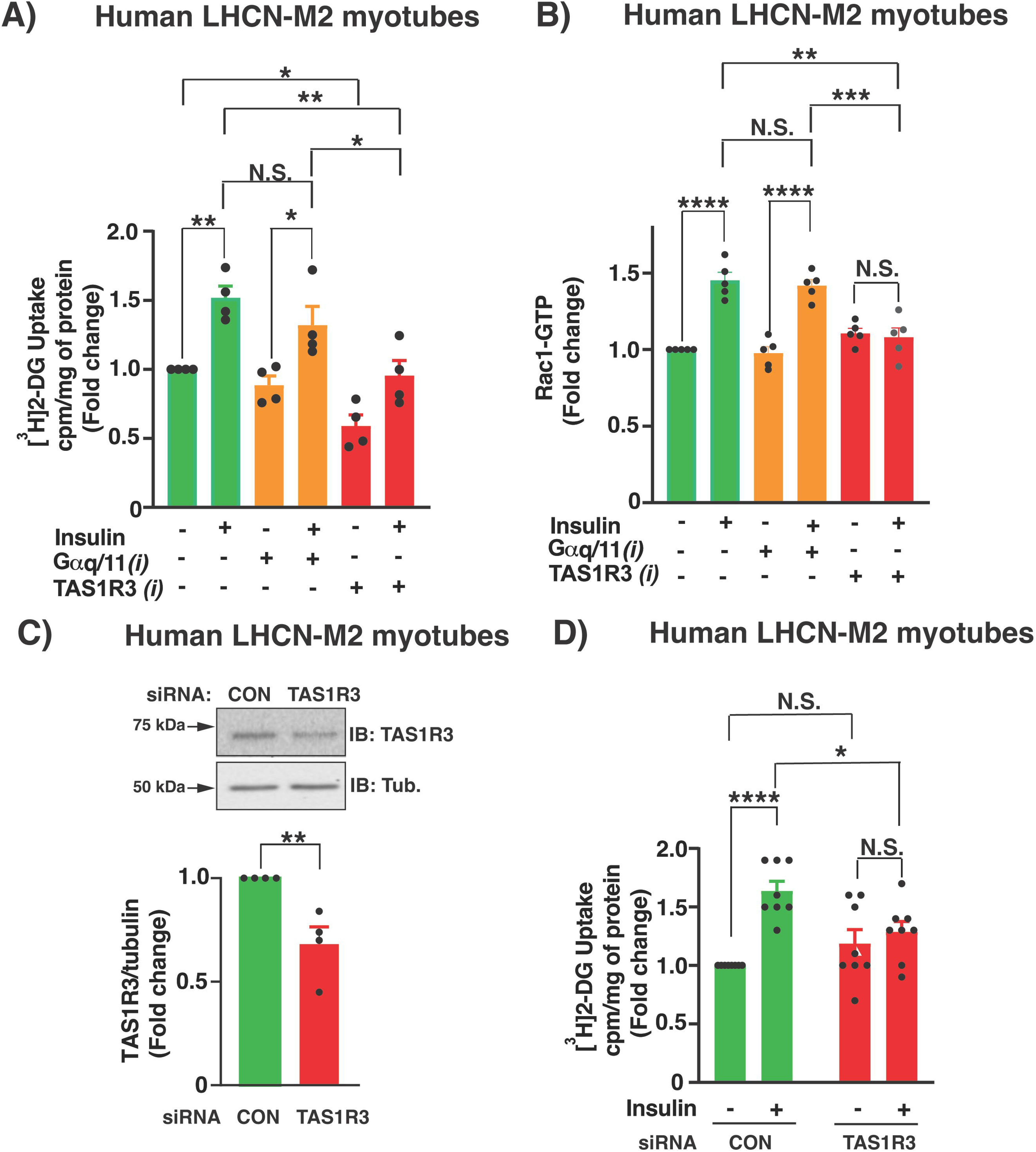
Lactisole inhibits insulin-stimulated glucose uptake and Rac1 activation in human LHCN-M2 myotubes. (A-B) Human LHCN-M2 myotubes were pre-treated with vehicle (DMSO) or inhibitor for 1 h and then stimulated with ± 100 nM insulin 5 minor 20 min for Rac1 activation and glucose uptake assay respectively. TAS1R3*(i)*; lactisole. Gαq/11(*i*); YM-25490. Assessment of [^3^H]2-DG uptake (**A**) and Rac1 activation (**B**). (**A**) Glucose uptake was normalized to protein content. Cpm: Counts per minute (n=4 independent experiments). (**B**) Rac1 activation values are relative to baseline Rac1-GTP levels (set to 1.0) detected with unstimulated basal insulin (n=5 independent experiments). **(C-D)** Human LHCN-M2 myotubes were pre-treated with siRNA (48 h) and then stimulated with ± 100 nM insulin (10 min). CON: control. (C) Impact of TAS1R3 siRNA knockdown. Top: Representative immunoblot (IB). Bottom: Densitometry analysis of TAS1R3 protein expression normalized to tubulin (Tub). Control-siRNA values were set to 1.0. N=4 independent experiments. (**D**) Effects of siRNA treatment on glucose uptake with insulin stimulation (n= 8 independent experiments). Glucose uptake was normalized to protein content. Cpm: Counts per minute. Values under basal unstimulated insulin conditions were set to 1.0. (**A-D**) Data expressed as mean ± SEM. * p < 0.05, ** p < 0.01, *** p < 0.001 and **** p < 0.0001. N.S: not significant.

Activation of the small GTPase Rac1 is required to mobilize GLUT4-laden vesicles to supply the GLUT4 protein at the sarcolemma to facilitate insulin-stimulated glucose uptake *via* a non-canonical insulin pathway (39). We tested the requirement for TAS1R3 in Rac1 activation in human LHCN-M2 myotubes by inhibiting TAS1R3 and comparing with Gαq/11 inhibition. While neither inhibitor altered Rac1-GTP levels during basal conditions (**Fig. 2B**), only TAS1R3 inhibition significantly reduced insulin-stimulated Rac1-GTP levels (**Fig. 2B**). We verified that indeed the decrease in insulin-stimulated glucose uptake in myotubes was TAS1R3 specific *via* TAS1R3 small interfering RNA (siRNA) knockdown (**Fig. 2C-D**). Thus, these results indicate a selective role for TAS1R3 in insulin-stimulated glucose uptake in human LHCN-M2 myotubes.

We next investigated the impact of TAS1R3 inhibition on canonical and non-canonical insulin signaling cascades (22–24, 40). Canonical insulin-signaling was assessed in LHCN-M2 myotubes, and TAS1R3 inhibition did not impact phosphorylation (activation) of canonical insulin cascade components IRS1 (Tyr^608^) and AKT (Ser^473^) (**Fig. 3A-B**). Non-canonical insulin signaling downstream of Rac1 involves Cofilin, a protein that oversees actin remodeling to translocate GLUT4-laden vesicles to the plasma membrane; Cofilin phosphorylation leads to depolymerization of actin filaments, which impedes GLUT4 vesicle translocation in skeletal muscle cells (41, 42). Vehicle-treated insulin-stimulated myotubes showed >50% decrease in p-Cofilin (relative to total cofilin protein); no effect on p-Cofilin was observed with TAS1R3*i*-treated insulin-stimulated myotubes (**Fig. 3C**). As observed above (**Fig. 2**), Gαq/11 inhibition did not impact the insulin signaling components (**Fig. 3A-C**). Taken together, our results suggest that TAS1R3 functions in non-canonical insulin signaling *via* Rac1 activation to promote insulin-stimulated glucose uptake in LHCN-M2 myotube.

**Figure 3.**
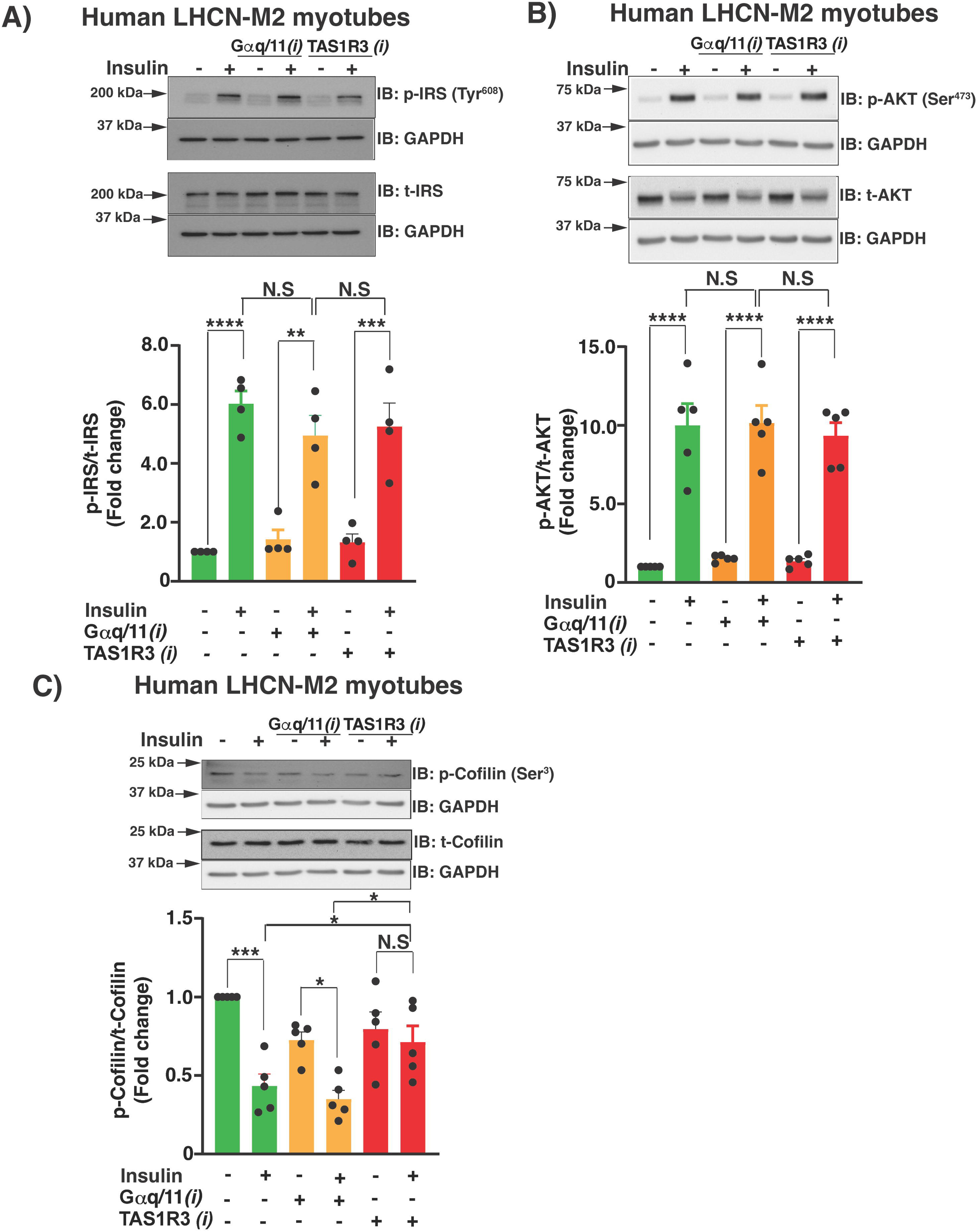
Lactisole fails to inhibit the activation of IRS and AKT but prevents the dephosphorylation of cofilin. **(A-C)** Human LHCN-M2 myotubes were pre-treated with vehicle (DMSO), TAS1R3*i* lactisole, or Gαq/11*i* YM-25490 for 1 h and were stimulated with ± 100 nM insulin (10 min). A representative immunoblot and densitometry analysis is shown for: **(A)** phospho-IRS (Tyr^608^) and total IRS (n=4 independent experiments), **(B)** phospho-AKT (Ser^473^) and total AKT (n=5 independent experiments), and **(C)** phospho-cofilin (Ser^3^) and total cofilin (n=5 independent experiments). Data expressed as mean ± SEM. * p < 0.05, ** p < 0.01, *** p < 0.001 and **** p < 0.0001. N.S: not significant.

## DISCUSSION

This study provides mechanistic insights into the role of TAS1R3-mediated monomeric GTPase signaling in skeletal muscle insulin-stimulated glucose uptake. For the first time, we showed that T2D significantly reduces TAS1R3 mRNA and protein levels in human skeletal muscle. Furthermore, we demonstrated that both direct pharmacological inhibition and siRNA knockdown of TAS1R3 contributes to decreased insulin-stimulated glucose uptake in human LHCN-M2 myotubes, consistent with the impaired glucose tolerance and insulin resistance phenotype observed in the TAS1R3 knockout mice (27). In this study, TAS1R3 function in skeletal muscle glucose uptake was attributed to the non-canonical Rac1 mediated insulin cascade for GLUT4 vesicle translocation to the sarcolemma, which is independent of Gαq/11 signaling. Furthermore, we observed TAS1R3 inhibition reduced glucose uptake under basal conditions (**Fig. 2A**), akin to Rac1 GTPase inhibitors on soleus muscle under basal conditions (40), TAS1R3 blockade of basal glucose, suggests that TAS1R3 supports basal glucose uptake, potentially through nutrient-sensing mechanisms intrinsic to skeletal muscle independently of insulin signaling. Collectively, the results highlight a role for TAS1R3 in skeletal muscle insulin-stimulated glucose uptake.

A key finding of this study was the essential role TAS1R3 plays as a GPCR target to initiate the non-insulin signaling cascade in human myotubes for insulin-stimulated glucose uptake *via* small GTPase Rac1 activation, which in turn promotes actin cytoskeletal remodeling *via* phospho-mediated inactivation of cofilin for GLUT4-vesicle translocation to the sarcolemma. Although we identified that TAS1R3 regulates this independently of Gαq/11 signaling, the precise trimeric GTPase that links TAS1R3 to downstream pathways remains to be elucidated. Among the numerous G-protein candidates, Gα14 stands out as a particularly promising mediator of skeletal muscle glucose uptake. This is supported by findings showing that FR900359, a selective inhibitor of Gαq/11/14 significantly impairs both GLUT4 translocation and glucose uptake in skeletal muscle cells (37) likely through an insulin-independent mechanism involving AMPK activation. In contrast, our experiments using YM-254890, a Gαq/11-specific inhibitor, failed to replicate these effects, suggesting that Gα14 may play a unique and critical role in this signaling axis. Nevertheless, the possibility that YM-254890 modulates AMPK activity cannot be entirely excluded, and further investigation is warranted to delineate its precise impact. Thus, future studies are required to determine the key G-protein candidate(s) downstream of TAS1R3 to complete the missing link between TAS1R3 activity and Rac1-GTP mediated GLUT4-vesicle translocation for insulin-stimulated glucose uptake in skeletal muscle cells in vitro.

These findings align with emerging evidence from TAS1R2 signaling, where glucose stimulation of this GPCR activates ERK1/2-dependent phosphorylation of PARP1, a major NAD consumer in skeletal muscle (28). Muscle-specific TAS1R2 knockout in mice suppressed PARP1 activity, elevated NAD levels, enhanced mitochondrial function, and improved endurance, positioning TAS1R2 as a peripheral energy sensor (28). Together, these studies underscore the broader role of GPCRs like TAS1R2 and TAS1R3 in regulating skeletal muscle metabolism through distinct but complementary pathways involving glucose sensing, cytoskeletal remodeling and NAD homeostasis. Future work is needed to identify the specific trimeric GTPase(s) downstream of TAS1R3 and to determine whether TAS1R3, like TAS1R2, contributes to mitochondrial health and endurance capacity.

We showed that TAS1R3 mRNA and protein levels were depleted in T2D skeletal muscle. Previously, TAS1R3 was shown to be involved in the musculoskeletal system, including myogenesis, amino acid sensing *via* mTOR activation to suppress autophagy (43) and osteoclastogenesis for bone remodeling and homeostasis (44). Importantly, the long-term secondary complication from prolonged T2D include dysfunctions across these aforementioned functions/pathways culminating in musculoskeletal comorbidities that include reduced bone quality, weakened muscle and impaired muscle repair. Conversely, the risks current T2D therapeutic interventions such as GLP-1 RAs pose on exacerbating these musculoskeletal comorbidities highlight the necessity of novel T2D therapeutics to mitigate and/or prevent these events. One limitation of the current study is the inability to determine the impact of skeletal muscle–specific TAS1R3 inactivation or depletion on whole-body glucose homeostasis. This highlights the need for cell type–specific, inducible TAS1R3 knockout animal models to enable in vivo investigations. Thus, future studies need to: [1] evaluate whether novel TAS1R3 agonists or genetic enrichment strategies can reverse skeletal muscle insulin resistance; and [2] develop skeletal muscle–specific, inducible TAS1R3 overexpression models to assess the potential benefits of TAS1R3 enrichment on primary insulin resistance and secondary muscle pathologies associated with T2D.

In conclusion, our findings provide novel insights into the functional activity of TAS1R3 as a modulator of insulin-stimulated glucose uptake *via* monomeric GTPase Rac1 (**Fig. 4**), independent of Gαq/11 signaling. Furthermore, we demonstrated that TAS1R3 is dysregulated in T2D skeletal muscle, suggesting a role in glucose homeostasis. Nevertheless, these emerging mechanistic insights establishes TAS1R3 as a future target to remediate skeletal muscle insulin resistance in prediabetes and T2D.

**Figure 4.**
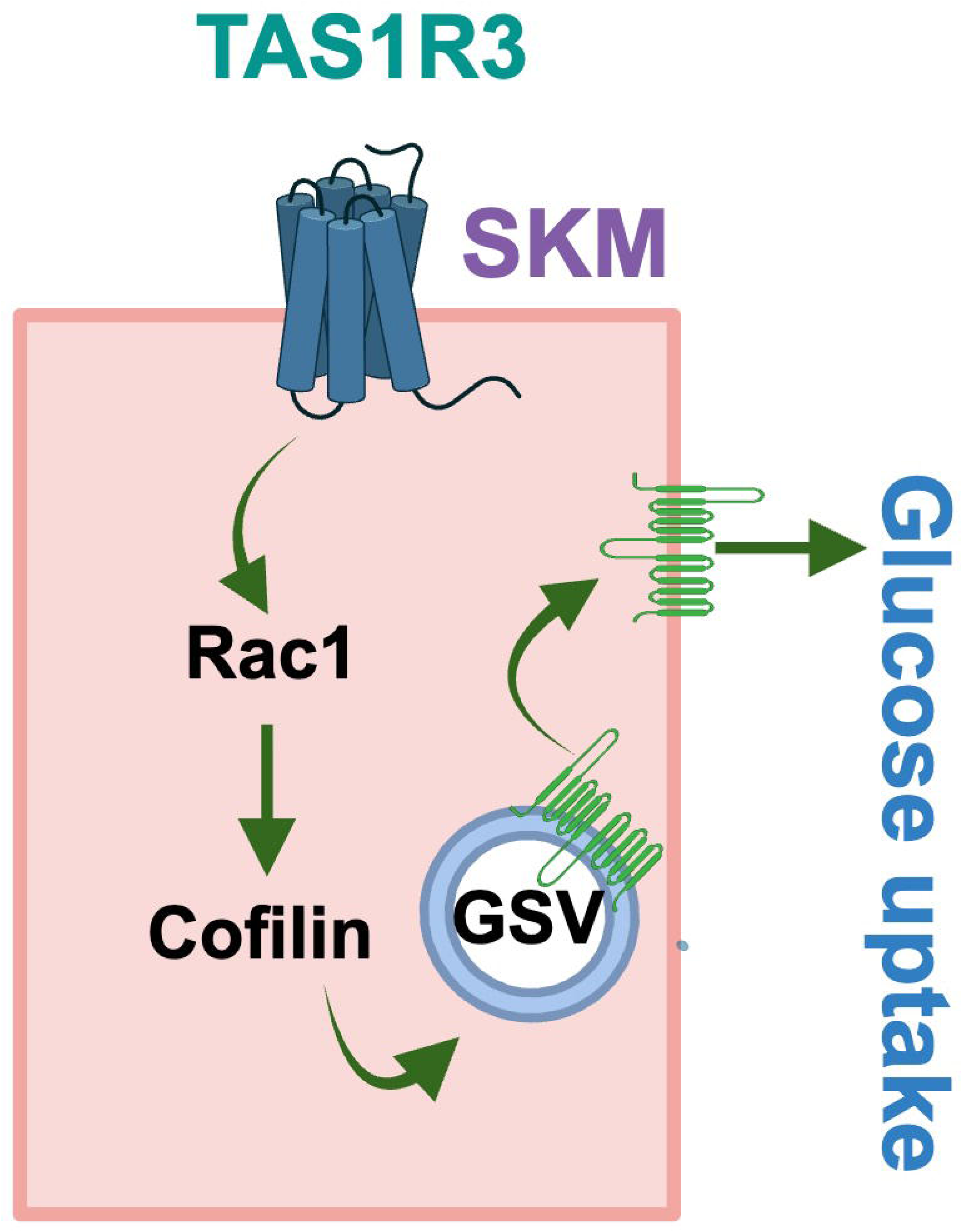
Schematic of TAS1R3 signaling in skeletal muscle. Activation of TAS1R3, a sweet taste receptor expressed in skeletal muscle, triggers a non-canonical signaling cascade that activates Rac1—a small GTPase essential for actin cytoskeletal remodeling. This sustained Rac1 activity prevents the insulin-driven dephosphorylation of Cofilin. Consequently, GLUT4 vesicle trafficking to the plasma membrane is impaired, leading to a marked reduction in glucose uptake.

## Materials and Methods

### LHCN-M2 Myoblast Cell Culture

For the physiological relevance of skeletal muscle glucose uptake, the immortalized human LHCN-M2 skeletal myoblast cell line was used (a generous gift from Dr. Melissa Bowerman, Keele University, UK). LHCN-M2 myoblasts were cultured in complete growth medium (4:1 ratio of 5 mM glucose DMEM (Thermo Fisher, Waltham, MA) and 5 mM glucose M199 medium (Thermo Fisher, Waltham, MA) supplemented with 15% [v/v] HI-FBS, 1% [v/v] antibiotic-antimycotic solution, 20 mM pH 7.4 HEPES buffer, 30 ng/mL Zn_2_SO_4_, 1.4 μg/mL vitamin B12 (Sigma-Aldrich, St. Louis, MO), 55 ng/mL dexamethasone (Sigma-Aldrich, St. Louis, MO), 2.5 ng/mL recombinant human hepatocyte growth factor (Sigma-Aldrich, St. Louis, MO), and 10 ng/mL recombinant human basic FGF (Biopioneer, San Diego, CA) on plastic cell culture dishes pre-coated with 1% [w/v] autoclaved porcine gelatin (Sigma-Aldrich, St. Louis, MO). At >70% confluence, LHCN-M2 myoblasts were cultured in complete differentiation medium composed of DMEM/M199 4:1, 2% [v/v] heat-inactivated horse serum (HI-HS) (Thermo Fisher, Waltham, MA), 1% [v/v] anti-biotic/anti-mycotic solution, 20 mM pH 7.4 HEPES buffer, 30 ng/mL Zn_2_SO_4_, and 1.4 μg/mL vitamin B12 for 6 days with fresh media changes every 48 h in 37°C and 5% CO_2_ incubator until multi-nucleated myotubes formed. For glucolipotoxicity (GLT)-induced diabetogenic stress, LHCN-M2 myotubes were exposed to 200 μM palmitate and 25 mM glucose for 16 h.

### Human skeletal muscle

Cadaveric non-diabetic and T2D skeletal leg muscles were purchased from the National Disease Research Interchange (NDRI, Philadelphia, PA). See **Table 1** for donor information. The samples were snap-frozen and kept at −80°C until mRNA and protein extraction were carried out.

### Quantitative PCR

Total RNA isolation from human donor samples and LHCN-M2 myotubes was performed with TriReagent (Millipore Sigma, St. Louis, MO, USA) as previously described (45) and from LHCN-M2 cell line with RNeasy Plus Mini Kit (Qiagen) according to manufacturer’s instructions. Gene expression was assessed using two-step reverse transcription (iScript^TM^ cDNA Synthesis Kit, Bio-Rad, Hercules, CA) and qPCR (iQ SYBR® Green Supermix, Bio-Rad, Hercules, CA). The cycle threshold data was converted to change-fold in expression by the “ΔΔCt” method.

### Immunoblot analysis

Whole protein lysates of cell cultures and primary human skeletal muscle biopsies were lysed with 1% NP-40 lysis buffer, resolved and underwent immunodetection as previously described (46). For detection of insulin signaling pathway, membranes were incubated with primary antibodies for IRS (Tyr^608^, 1: 1,000, Millipore-Sigma, St. Louis, MO), AKT (Ser^473^, 1:1,000, Cell Signaling Technology, Danvers, MA) (46) and cofilin (Ser^3^, 1:1,000, Santa Cruz Biotechnology, Inc. Dallas, TX) (41). As a control for equivalent protein-loading, IRS (1:1000, Cell Signaling Technology, Danvers, MA), AKT (1:1,000, Cell Signaling Technology, Danvers, MA) and Cofillin (1:1,000, Cell Signaling Technology, Danvers, MA) and GAPDH (1:50 000, Invitrogen, Waltham, MA) antibodies were used.

### LHCN-M2 2-Deoxyglucose Uptake

For the pharmacological analysis, LHCN-M2 myotubes were subjected to two washes with an oxygenated FCB buffer devoid of serum and glucose. This buffer composed of 125 mM NaCl, 5 mM KCl, 1.8 mM CaCl2, 2.6 mM MgSO4, 25 mM HEPES, 2 mM pyruvate, and supplemented with 2% (wt/vol) bovine serum albumin (BSA). Subsequently, the myotubes were treated with either vehicle (0.1% v/v DMSO) or 1 mM lactisole to specifically inhibit TAS1R3, or 10 µM YM-25490, a selective antagonist for Gαq/11 signaling. This treatment was conducted in the aforementioned FCB buffer for 1h. Following drug exposure, insulin stimulation was performed using 100 nM insulin for 20 min. The myotubes were then assessed for 2-deoxyglucose (2-DG) uptake, following methodologies previously established for L6.GLUT4myc myotubes (34, 46, 47).

### Small Interfering RNA Transfection

For TAS1R3 knockdown studies, LHCN-M2 myotubes at differentiation day (D) 4 were transfected with 50 nM TAS1R3-targeting siRNA (siTAS1R3) sequence or non-specific targeting siRNA control in a drop wise fashion using RNAiMax and Opti-MEM overnight as previously described (47). These myotubes underwent a total of 48 h transfection prior to 2-DG uptake protocol (see above).

### G-LISA for Rac1 activation assay

Rac1 activation was measured in the cell lysate from LHCN-M2 myotubes using the commercially available G-LISA Rac1 Activation Assay from Cytoskeleton Inc (Denver, CO). In brief, the LHCN-M2 myotubes were washed twice with PBS and then serum-and glucose-starved in oxygenated FCB buffer for one hour. During this time, the myotubes were treated with either vehicle or 1 mM lactisole, or 10 µM YM-254890 to induce pharmacological inhibition of TAS1R3 (lactisole) or Gq/11 (YM-254890). This was followed by a 5min stimulation with 100 nM insulin, and comparisons were made to basal controls with and without drug treatment. Rac1 activation was measured in the supernatant using the commercially available G-LISA Rac1 Activation Assay from Cytoskeleton Inc (Cat # BK 128) as described previously (48). Intensity of the color formed from the chromogenic substrate was quantified by colorimetry using a BioTek Synergy HTX Multi-Mode Plate Reader.

### Insulin signaling analysis

For the detection of insulin signaling pathway, LHCN-M2 myotubes underwent 1h incubation in serum- and glucose-starved oxygenated FCB buffer with either vehicle, 1 mM lactisole or 10 μM YM-254890 prior to 100 nM insulin stimulation for 10 min as described for L6.GLUT4myc myotubes (46).

### Statistics

Data are presented as mean ± SEM and *n* values are indicated in the figures. Differences between two groups were assessed using Student’s t-test. Statistically significant differences among multiple groups were evaluated using one-way or two-way ANOVA followed by Bonferroni post hoc test. The threshold for statistical significance was set at p < 0.05 and all the data were analyzed using GraphPad Prism software, version 8.3.0. Statistical significance is indicated in the figure legends.

### Data and Resource Availability

The data sets generated during and/or analyzed during the current study are available from the corresponding author upon reasonable request. No applicable resources were generated during this study.

### ARTICLE INFORMATION

## Acknowledgements

The authors would also like to thank Pablo Garcia and Erika McCown for their technical assistance. We thank Dr. Chathurani S. Jayasena (COH) for providing critical feedback and editing the manuscript. We appreciate the valuable input and thoughtful suggestions provided by Drs. Patrick Fueger and Ke Ma, which contributed significantly to improving the clarity and impact of this work.

## Funding

This work was supported by the National Institutes of Health (DK067912 and DK112917 to D.C.T, and DK102233 to D.C.T and R.V) and The Arthur Riggs Diabetes and Metabolism Research Institute (COH) Pilot program (R.V) and an American Heart Association Career Development Award to RB (25CDA1454374).

## Duality of Interest

No potential conflicts of interest relevant to this article were reported.

## Author Contributions

R.V conceptualized the project. D.C.T and R.V acquired the funding. J.H, K.E.M., R.V, and R.B performed the experiments and 2-DG uptake analysis. D.C.T, J.H, R.B and R.V defined the methodology used. D.C.T supervised the project. R.V wrote the original draft of this manuscript. D.C.T, J.H, R.B and R.V. were involved in the data analysis, discussion and editing of the manuscript. All authors approved the final version of the manuscript. R.V is the guarantor of this work and, as such, had full access to all the data in the study and takes responsibility for the integrity of the data and the accuracy of the data analysis.

## Notes

### Competing Interest Statement

The authors have declared no competing interest.

### Summary of Updates

Our new findings demonstrate that GLT-induced stress (glucolipotoxicity) significantly reduces glucose uptake, consistent with an insulin resistance model. Under these conditions, TAS1R3 expression is markedly downregulated at the mRNA level, suggesting that impaired TAS1R3 signaling may contribute to the observed metabolic dysfunction. Furthermore, TAS1R3 inhibition modulates non-canonical Rac1-mediated signaling pathways, leading to decreased glucose uptake. These results expand the functional role of TAS1R3 beyond sweet taste perception, linking its inhibition to defective metabolic responses through Rac1-dependent mechanisms. Collectively, these findings highlights the translational potential of TAS1R3 as a therapeutic target for restoring glucose homeostasis.

